# Functional loss of a non-canonical BCOR-PRC1.1 complex accelerates SHH-driven medulloblastoma formation

**DOI:** 10.1101/2020.02.06.938035

**Authors:** Lena M. Kutscher, Konstantin Okonechnikov, Nadja V. Batora, Jessica Clark, Patricia B. G. Silva, Mikaella Vouri, Sjoerd van Rijn, Laura Sieber, Britta Statz, Micah D. Gearhart, Norman Mack, Brent A. Orr, Andrey Korshunov, Audrey L. Mercier, Olivier Ayrault, Marcel Kool, Vivian J. Bardwell, Stefan M. Pfister, Paul A. Northcott, Daisuke Kawauchi

**Author notes:** National Institute of Neuroscience, NCNP, Tokyo, Japan, 187-0031. Co-corresponding authors, DK; PAN. These authors contributed equally to this work (Lena M. Kutscher, Konstantin Okonechnikov, Nadja V. Batora).

## Abstract

Medulloblastoma is a childhood brain tumor arising from the developing cerebellum. In Sonic Hedgehog (SHH)-subgroup medulloblastoma, aberrant activation of SHH signaling causes increased proliferation of granule neuron progenitors (GNPs) and predisposes these cells to tumorigenesis. A second, cooperating genetic hit is often required to push these hyperplastic cells to malignancy and confer mutation-specific characteristics associated with oncogenic signaling. Somatic loss-of-function mutations of the transcriptional co-repressor *BCOR* are recurrent and highly enriched in SHH-medulloblastoma. To investigate *BCOR* as a putative tumor suppressor, we used a germline genetically engineered mouse model to delete exons 9/10 of *Bcor* (*Bcor*^*ΔE9-10*^) in GNPs during development. This leads to reduced expression of C-terminally truncated BCOR (BCOR^ΔE9-10^). While *Bcor*^*ΔE9-10*^ alone did not promote tumorigenesis or affect GNP differentiation, *Bcor*^*ΔE9-10*^ combined with loss of the SHH-receptor gene *Ptch1* resulted in highly penetrant medulloblastomas. In *Ptch1+/-;Bcor*^*ΔE9-10*^ tumors, the growth factor gene *Igf2* was aberrantly upregulated, and ectopic *Igf2* overexpression was sufficient to drive tumorigenesis in *Ptch1*+/- GNPs. BCOR directly regulates *Igf2*, likely through the PRC1.1 complex; the repressive histone mark H2AK119Ub is decreased at the *Igf2* promoter in *Ptch1+/-;Bcor*^*ΔE9-10*^ tumors. Overall, our data suggests that BCOR-PRC1.1 disruption leads to *Igf2* overexpression, which transforms preneoplastic cells to malignant tumors.

## INTRODUCTION

Pediatric cancer is the number one cause of disease-related death in children, and brain tumors are the most common pediatric solid tumor (Udaka and Packer 2018). Medulloblastoma is an embryonic cerebellar tumor comprised of at least four biologically and clinically distinct molecular subgroups (Northcott et al. 2012). One subgroup, Sonic Hedgehog (SHH)-medulloblastoma, arises through genetic alterations resulting in activation of the SHH signaling pathway, including mutations in the SHH receptor gene *PTCH1*, activating mutations in the transmembrane protein-coding gene *Smoothened* (*SMO*), mutations in the negative regulator gene *SUFU*, or amplification of the transcription factor genes *GLI2* or *MYCN* (Johnson et al. 1996; Hahn et al. 1996; Goodrich et al. 1997; Kenney et al. 2003; Ayrault et al. 2010; Kool et al. 2014).

While constitutive activation of SHH signaling is required for SHH-medulloblastoma formation, studies in mice have demonstrated that additional genetic hits are required for the malignant transformation of cerebellar granule neuron progenitors (GNPs) in *Ptch1* heterozygous animals (Tamayo-Orrego and Charron 2019; Oliver et al. 2005; Kessler et al. 2009; Tamayo-Orrego et al. 2016). Depending on the nature of these mutations, the molecular and cellular properties of the tumor may change (Vogelstein and Kinzler 1993). More importantly, understanding how these cooperating mutations contribute to malignancy may reveal new therapeutic opportunities for affected patients.

We and others previously identified recurrent inactivating mutations and insertions/deletions (indels) targeting the transcriptional co-repressor gene *BCOR* in SHH-medulloblastoma (Jones et al. 2012; Pugh et al. 2012; Robinson et al. 2012; Kool et al. 2014; Northcott et al. 2017), but the mechanism(s) underlying *BCOR-*associated medulloblastoma remain unclear. *BCOR* aberrations are implicated in a variety of pediatric cancers, including acute myeloid leukemia, retinoblastoma, sarcomas, glioblastomas, and CNS high-grade neuroepithelial tumor with *BCOR* alteration (CNS-HGNET-BCOR) (Mackay et al. 2017; Astolfi et al. 2015; Sturm et al. 2016; Kooi et al. 2016; Grossmann et al. 2011; Santiago et al. 2017). Altogether, deregulation of *BCOR* is implicated in at least 18 different tumor histotypes of pediatric cancer (Astolfi et al. 2019), demonstrating the urgency in understanding *BCOR* function in both normal and tumorigenic cells.

BCOR was originally identified based on its interaction with the zinc finger transcription factor BCL6 and subsequently shown to be a component of a non-canonical Polycomb Repressive Complex (PRC1.1) (Huynh et al. 2000; Gearhart et al. 2006; Cao et al. 2016; Gao et al. 2012). BCL6 can recruit BCOR to DNA via the BCL6-interacting motif found in the N-terminal half of BCOR (Fig. 1A) (Ghetu et al. 2008). The C-terminus contains non-ankyrin repeats, ankyrin repeats, and the PUFD domain, which binds the Polycomb group protein PCGF1 (Fig. 1A) (Junco et al. 2013). These regions and their interacting partners are conserved in BCOR in mice (Wamstad and Bardwell 2007; Huynh et al. 2000; Wamstad et al. 2008).

**Figure 1.**
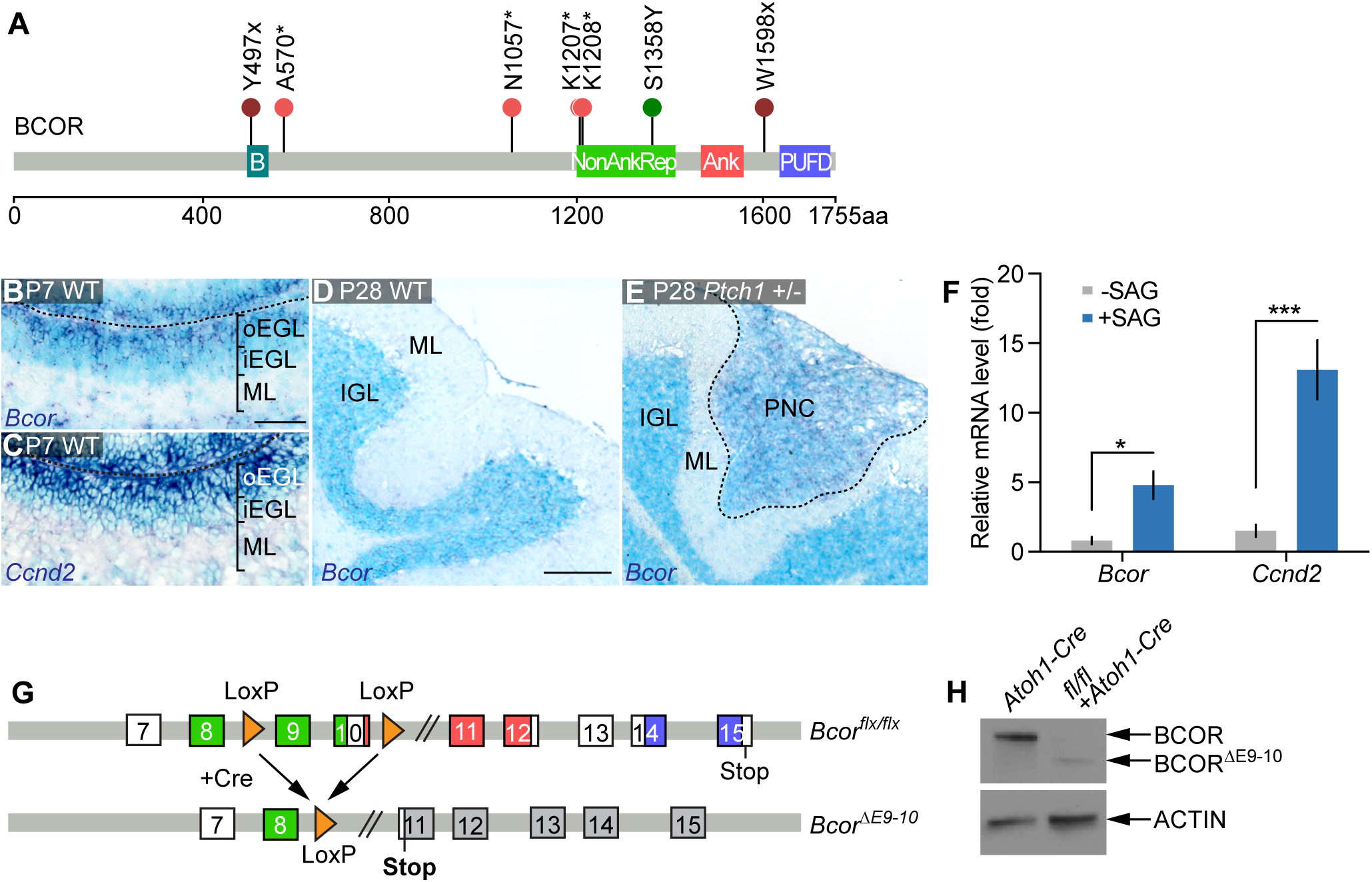
Conditional *Bcor* mouse model is an appropriate model for human *BCOR* mutations. **(A)** BCOR mutations identified in pediatric medulloblastoma samples. B, BCL6 binding domain; NonAnkRep, Non-Ankyrin Repeats; Ank, Ankyrin repeats; PUFD, PCGF Ub-like fold discriminator domain; maroon lollipop, nonsense mutation; pink lollipop, frame shift mutation; green lollipop, point mutation. (**B**,**C**) *in situ* hybridization (dark blue) of antisense (**B**) *Bcor* probe and (**C**) *Ccnd2* probe at P7 in Bl6N wild type (WT) mice. Scale bar, 50 μm. oEGL, outer external granule layer. iEGL, inner external granule layer. ML, molecular layer. (**D**,**E**) *in situ* hybridization of antisense *Bcor* probe at P28 in (**D**) WT and in (**E**) *Ptch1*+/- mice. Scale bar, 200 μm. IGL, inner granule layer. ML, molecular layer. PNC, preneoplastic cells. (**F**) Relative mRNA expression of *Bcor* and *Ccnd2* (quantitative PCR) of P7 granule neuron progenitors cultured in the absence (gray) or presence (blue) of Smoothened agonist (SAG). *, p<0.05, ***, p<0.001, Student’s T test. Bars represent mean +/- SEM. N=6. (**G**) Location of LoxP sites in *Bcor*. Expression of Cre removes exons 9 and 10, and leads to an early translation stop. Color scheme for *Bcor* domains same as in (**A**). (**H**) anti-BCOR western blot of purified granule neuron progenitors in wild type (*Atoh1-Cre only*) cells and in *Bcor*^*floxed/floxed*^ + *Atoh1-Cre* (*Bcor*^*ΔE9-10*^) cells. Anti-ACTIN, loading control. See also Supplemental Figure S1.

BCL6 has been previously implicated in cerebellar development in mice (Tiberi et al. 2014). Overexpression of human *BCL6* suppresses medulloblastoma formation *in vivo* in mice, perhaps through interaction with BCOR at *Gli1/Gli2* promoters (Tiberi et al. 2014), although recurrent *BCL6* mutations have not yet been identified in SHH-medulloblastoma patient samples. While the interaction between BCOR and BCL6 may be one mode of BCOR-mediated tumor suppression, *BCOR* mutations seen in SHH-medulloblastoma frequently truncate the C-terminal PUFD domain (Fig. 1A) (Northcott et al. 2017). This truncation may lead to an inability to recruit PCGF1, which together with RING1B ubiquitinates histone H2A at lysine 119 (H2AK119Ub) to repress transcription of target genes (Gao et al. 2012; Gearhart et al. 2006; Aranda et al. 2015). Therefore, we set out to determine whether BCOR function within the PRC1.1 complex is involved in tumor suppression in SHH-medulloblastoma.

Here we show that genetic ablation of exons 9 and 10 of *Bcor* (*Bcor*^*ΔE9-10*^) in developing GNPs does not disrupt granule neuron differentiation in mice. Instead, *Bcor*^*ΔE9-10*^ cooperates with inactivation of *Ptch1* to potentiate aggressive medulloblastoma formation. BCOR occupies the *Igf2* promoter region in *Ptch1+/-* tumors, and chromatin occupancy is greatly reduced in *Ptch1+/-;Bcor*^*ΔE9-10*^ tumors. This reduction leads to aberrant upregulation of *Igf2* in these tumors to drive tumorigenesis. Additionally, BCOR^ΔE9-10^ no longer binds RING1B, the catalytic subunit of the PRC1.1 complex, which leads to a global reduction of the repressive histone mark H2AK119Ub, and a reduction of this mark specifically at the *Igf2* promoter, compared to *Ptch1+/-* tumors. Our work demonstrates that loss of the C-terminus of BCOR disrupts PRC1.1 complex function to drive SHH-medulloblastoma formation.

## RESULTS

### BCOR is recurrently co-mutated with members of the SHH pathway in pediatric SHH-medulloblastoma

We and others previously identified *BCOR* as recurrently mutated or deleted in ∼10% of pediatric SHH-medulloblastomas (Jones et al. 2012; Pugh et al. 2012; Robinson et al. 2012; Kool et al. 2014; Northcott et al. 2017). We mapped the identified mutations to the *BCOR* coding sequence (Fig. 1A) and determined that the C-terminal PUFD domain is inferred to be absent because of the introduction of premature STOP codons in 6/7 cases (Table 1). In the majority of cases (6/7), *BCOR* is co-mutated with members of the SHH pathway, including *PTCH1* (4/6), *GLI2* (1/6), *SMO* (1/6) or *SUFU* (1/6). *BCOR* is an X-linked gene, and the majority of affected patients were male (6/7). Given its potential as a cooperating mutation in SHH-medulloblastoma formation, we investigated the functional relevance of these mutations during normal cerebellar development and tumorigenesis.

**Table 1.**
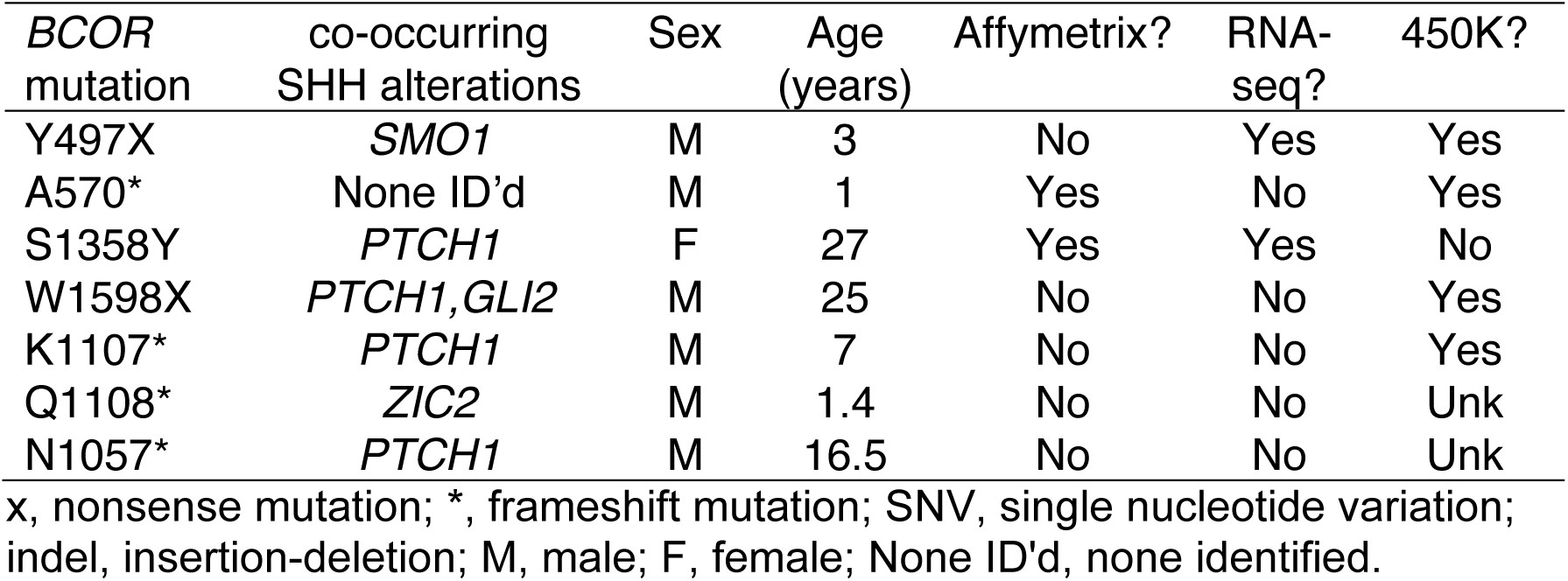
*BCOR* mutations in SHH-medulloblastoma patient samples.

| *BCOR* mutation | co-occurring SHH alterations | Sex | Age (years) | Affymetrix? | RNA-seq? | 450K? |
| --- | --- | --- | --- | --- | --- | --- |
| Y497X | *SMO1* | M | 3 | No | Yes | Yes |
| A570* | None ID'd | M | 1 | Yes | No | Yes |
| S1358Y | *PTCH1* | F | 27 | Yes | Yes | No |
| W1598X | *PTCH1, GLI2* | M | 25 | No | No | Yes |
| K1107* | *PTCH1* | M | 7 | No | No | Yes |
| Q1108* | *ZIC2* | M | 1.4 | No | No | Unk |
| N1057* | *PTCH1* | M | 16.5 | No | No | Unk |

x, nonsense mutation; \*, frameshift mutation; SNV, single nucleotide variation; indel, insertion-deletion; M, male; F, female; None ID'd, none identified.

### Bcor is expressed in proliferating granule neuron progenitors (GNPs) in mice

We first determined whether *Bcor* is normally expressed in developing cerebellar GNPs in mice, the cell-of-origin for SHH-medulloblastoma (Yang et al. 2008; Schüller et al. 2008; Hovestadt et al. 2019; Vladoiu et al. 2019). Using RNA *in situ* hybridization (ISH), we found that *Bcor* is expressed in the outer external granule layer of the P7 cerebellum (Fig. 1B), similar to actively cycling cells (*Ccnd2*, Fig. 1C). After cerebellar development is complete, however, *Bcor* is no longer expressed (P28, Fig. 1D).

In mice heterozygous for the SHH receptor gene *Ptch1* (written as *Ptch1*+/- hereafter), preneoplastic lesions form in 85% of mice by P21, but the majority regress by P42 (Corcoran et al. 2008; Kessler et al. 2009). We found that *Bcor* is expressed in these preneoplastic lesions but not in adjacent normal tissue (Fig. 1E).

To validate the *in vivo* expression pattern, we examined *Bcor* RNA levels in GNPs cultured with or without a Smoothened agonist (SAG) *in vitro. Bcor* expression is approximately 6x higher in proliferating GNPs (with SAG) compared to differentiated cells (without SAG, Fig. 1F, *Ccnd2* used as positive control). Together, the expression pattern of *Bcor* in normal and preneoplastic GNPs suggests that our mouse model is appropriate to study *Bcor* function in SHH-medulloblastoma.

### Bcor^ΔE9-10^ mouse model does not noticeably alter BCL6-related cerebellar functions, including GNP differentiation or migration

The mutations we identified in human tumors suggest the C-terminal domain of BCOR may be important for its tumor suppressive function (Fig. 1A). To investigate the functional relevance of these mutations during GNP development and tumorigenesis, we used germline genetically engineered mouse strains to generate Cre inducible excision of *Bcor* exons 9 and 10 in *Atoh1*-positive GNPs (*Bcor*^*ΔE9-10*^, Fig. 1G, see Methods for details (Tara et al. 2018)). At the protein level, this excision results in reduced expression of a truncated BCOR protein (Fig. 1H).

BCOR and its N-terminal interaction with BCL6 have been implicated in SHH-medulloblastoma (Tiberi et al. 2014). We sought to determine whether BCL6-related functions were disrupted in our mouse model. First, to investigate whether BCOR^ΔE9-10^ interacts with BCL6, we overexpressed HA-tagged full-length BCOR or HA-tagged BCOR^ΔE9-10^ with BCL6 in HEK293T cells and performed co-immunoprecipitation studies with the bulk protein extract. We found that BCOR^ΔE9-10^ interacts with BCL6 *in vitro* (Supplemental Fig. S1A), suggesting that the BCL6 interacting motif is still functional.

Next, we examined whether any BCL6-related BCOR functions may be affected *in vivo* in the *Bcor*^*ΔE9-10*^ mouse model. *Bcl6* is required for timely GNP differentiation (Tiberi et al. 2014), so we tested whether differentiation is affected in *Bcor*^*ΔE9-10*^ GNPs. In contrast to *Bcl6* knockout animals (Tiberi et al. 2014), we found that *Bcor*^*ΔE9-10*^ mice do not exhibit defects in GNP differentiation or migration (Supplemental Fig. S1B-K), further suggesting that previously reported BCL6-related functions in the cerebellum are largely unaffected in our mouse model. We found no differences between wild-type and *Bcor*^*ΔE9-10*^ animals in the total number of GNP cells at P7 (PAX6-positive, Supplemental Fig. S1B-D) or proliferating cells (Ki67-positive, Supplemental Fig. S1E-G). We also found no differences in proliferation or cell cycle exit at P8 after a single dose of EdU at P7 (Supplemental Fig. S1H-M). We examined migration of granule neuron cells at P28 following a single EdU pulse at P7 and found no differences between wild-type and *Bcor*^*ΔE9-10*^ animals (Supplemental Fig. S1N-R).

To further confirm that BCL6-dependent roles of BCOR are unaffected in *Bcor*^*ΔE9-10*^ GNPs, we examined expression of known BCL6/BCOR-regulated genes (Tiberi et al. 2014). Loss of *Bcl6* in cerebellar cells resulted in an increase in *Gli1* and *Gli2* expression, and both BCOR and BCL6 were bound at these promoter regions (Tiberi et al. 2014). To determine whether BCOR^ΔE9-10^ affects expression of these and other SHH-pathway related genes in GNPs, we isolated total RNA from wild-type and *Bcor*^*ΔE9-10*^ GNPs and evaluated target gene expression using qPCR. In contrast to *Bcl6* loss (Tiberi et al. 2014), loss of the region encoding the C-terminal domain of BCOR does not activate SHH signaling in P7 GNPs, including expression of *Gli1, Gli2, Ccnd1, Ptch1*, or *Mycn* (Supplemental Fig. S1S), suggesting that BCOR^ΔE9-10^ does not influence BCL6-related SHH signaling repression in GNPs. Taken together, our *Bcor*^*ΔE9-10*^ mouse model should allow us to assess the role of BCOR, and in particular the involvement of the C-terminus and PRC1.1, in medulloblastoma formation.

### Bcor^ΔE9-10^ significantly reduces latency and increases penetrance of Ptch1-associated medulloblastoma

Next, we examined how *Bcor*^*ΔE9-10*^ affects tumor formation in conjunction with *Ptch1* inactivation. Of note, deletion of the region encoding the C-terminal domain of BCOR in GNPs by itself does not promote medulloblastoma (Fig. 2A). Similar to previous studies (Goodrich et al. 1997), heterozygous mutations in *Ptch1* resulted in spontaneous medulloblastoma formation in 35% of animals, with a median latency of 179 days (Fig. 2A). In these mice, the second copy of *Ptch1* is mutated, with additional secondary mutations in other genes (Tamayo-Orrego et al. 2016). Combining *Bcor*^*ΔE9-10*^ with *Ptch1* mutations resulted in tumorigenesis with 100% penetrance and a median latency of 75 days (Fig. 2A, p<0.0001, Log-rank (Mantel-Cox) test). There was a histological switch from classic in *Ptch1*+/- tumors (N = 3/3, Fig. 2B) to the more aggressive histological subtype, large-cell anaplastic (LCA), in *Ptch1+/-;Bcor*^*ΔE9-10*^ tumors (N = 3/3, Fig. 2C). To confirm that *Ptch1+/-*;*Bcor*^*ΔE9-10*^ tumor cells had higher tumorigenic potential, we transplanted 8×10^5^ cells from either *Ptch1+/-* tumors or *Ptch1*+/-;*Bcor*^*ΔE9-10*^ tumors into the cerebella of immunodeficient mice. *Ptch1+/-;Bcor*^*ΔE9-10*^ tumor cells reestablished tumors considerably faster than *Ptch1+/-* tumor cells (Fig. 2D, median latency 26 days compared to 150 days, p<0.0001, Log-rank (Mantel-Cox) test).

**Figure 2.**
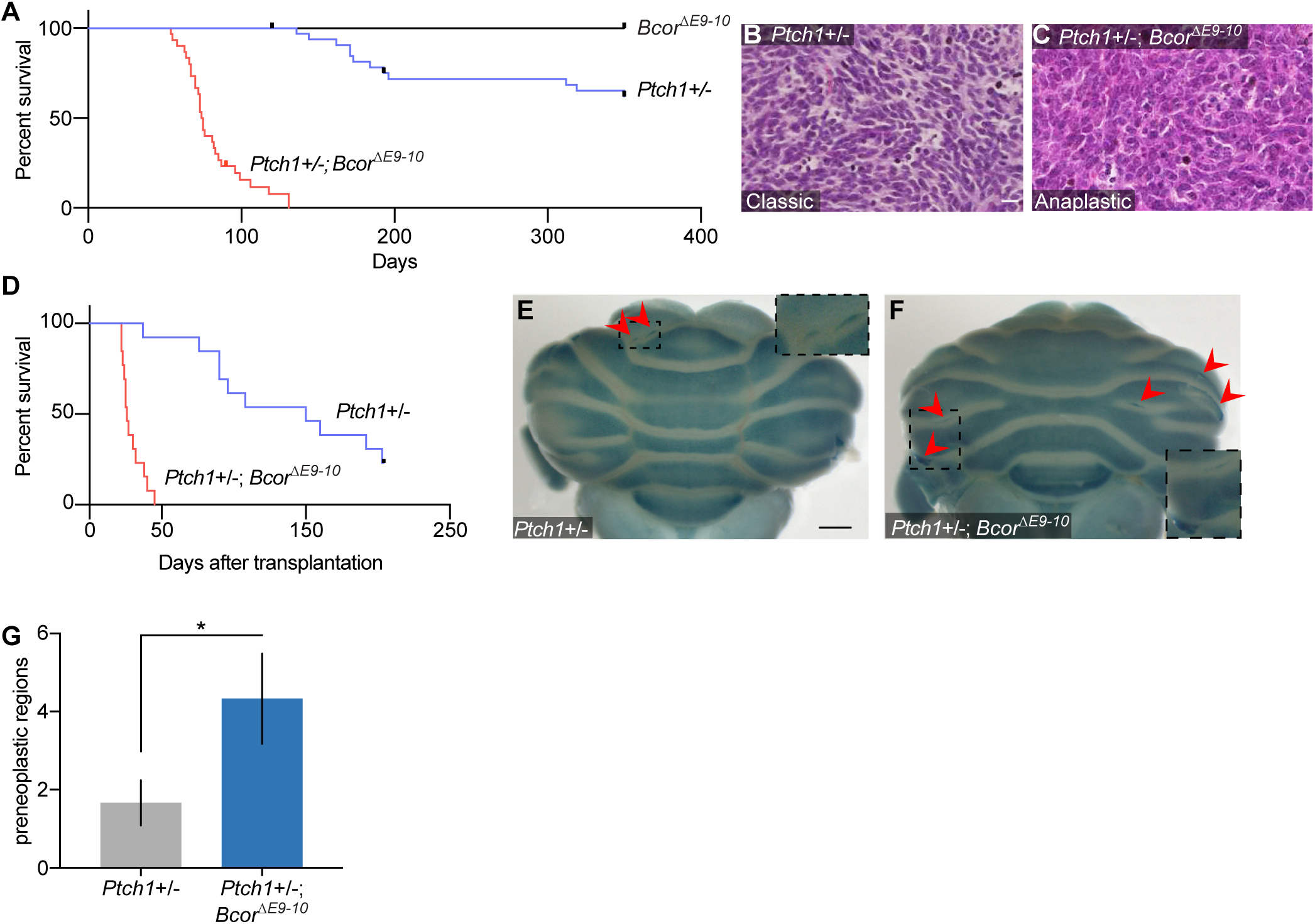
Mutations in *Bcor* significantly reduce latency and increase penetrance in *Ptch1*-driven medulloblastoma. (**A**) Survival curve of *Bcor*^*ΔE9-10*^ (black, N=10), *Ptch1*+/- (blue, N=34) and *Ptch1*+/-;*Bcor*^*ΔE9-10*^ (red, N=30) animals. (**B**,**C**) Representative H&E staining of (**B**) *Ptch1*+/- tumor and (**C**) *Ptch1*+/-;*Bcor*^*ΔE9-10*^ tumor. (**D**) Survival curve of immunodeficient animals transplanted with 1×10^6^ cells from *Ptch1*+/- (blue, N=13) and *Ptch1*+/-;*Bcor*^*ΔE9-10*^ (red, N=13) tumors. (**E**,**F**) Representative examples of beta-galactosidase staining (LacZ present in *Ptch1* locus) of P28 (**E**) *Ptch1*+/- cerebellum and (**F**) *Ptch1*+/-;*Bcor*^*ΔE9-10*^ cerebellum. Red arrows, preneoplastic lesions. Inset, magnified view of preneoplastic lesion. Scale bar, 1000μm. (**G**) Quantification of the number of preneoplastic lesions in (**E**,**F**). *, p<0.05, Student’s T test. Bars represent mean +/- standard deviation. N=3 cerebella per genotype.

To determine whether *Bcor*^*ΔE9-10*^ drives early transformation events, we next examined the number of preneoplastic lesions in the cerebella of P28 mice. *Ptch1*+/- animals carry a *lacZ* gene that disrupts one copy of the *Ptch1* locus, and preneoplastic cell clusters, where the *Ptch1* promoter is active, can be identified by beta-galactosidase staining (Fig. 2E,F, Goodrich et al. 1997; Oliver et al. 2005). We determined that *Ptch1+/-;Bcor*^*ΔE9-10*^ cerebella have twice the number of these regions compared to *Ptch1*+/- cerebella alone (Fig. 2G, N=3 cerebella per genotype, p=0.0232, Student’s t-test). This result suggests that a greater proportion of GNPs continue to proliferate in the absence of full-length *Bcor* and *Ptch1*, increasing the likelihood of malignant transformation. Intriguingly, loss of *Ptch1* together with *Bcor*^*ΔE9-10*^ results in a decrease of SHH-related gene expression compared to *Ptch1* alone (P7 GNPs, Supplemental Fig. S1S), suggesting deregulation of an alternative pathway(s) may be required to promote malignant transformation.

### Igf2 is upregulated in mouse and human BCOR-associated medulloblastomas

Based on our qPCR data (Supplemental Fig. S1S), SHH pathway genes are likely not the only drivers in *Ptch1+/-;Bcor*^*ΔE9-10*^ tumors. To identify alternative signaling pathways or genes that may contribute to tumorigenesis in this context, we used RNA-sequencing to identify misregulated genes in *Ptch1+/-;Bcor*^*ΔE9-10*^ tumors compared to *Ptch1+/-* tumors. Unsupervised clustering of transcriptome profiles demonstrated a clear difference between the tumor models (Supplemental Fig. S2A) and differential expression analysis revealed possible drivers (Supplemental Table S1). Among the 448 significantly upregulated genes, we identified the growth factor *Igf2* as upregulated ∼20-fold in *Ptch1+/-;Bcor*^*ΔE9-10*^ tumors compared to *Ptch1*+/- tumors (Fig. 3A, Log2FC: ∼4.4, adj. p = 5.26e-30).

**Figure 3.**
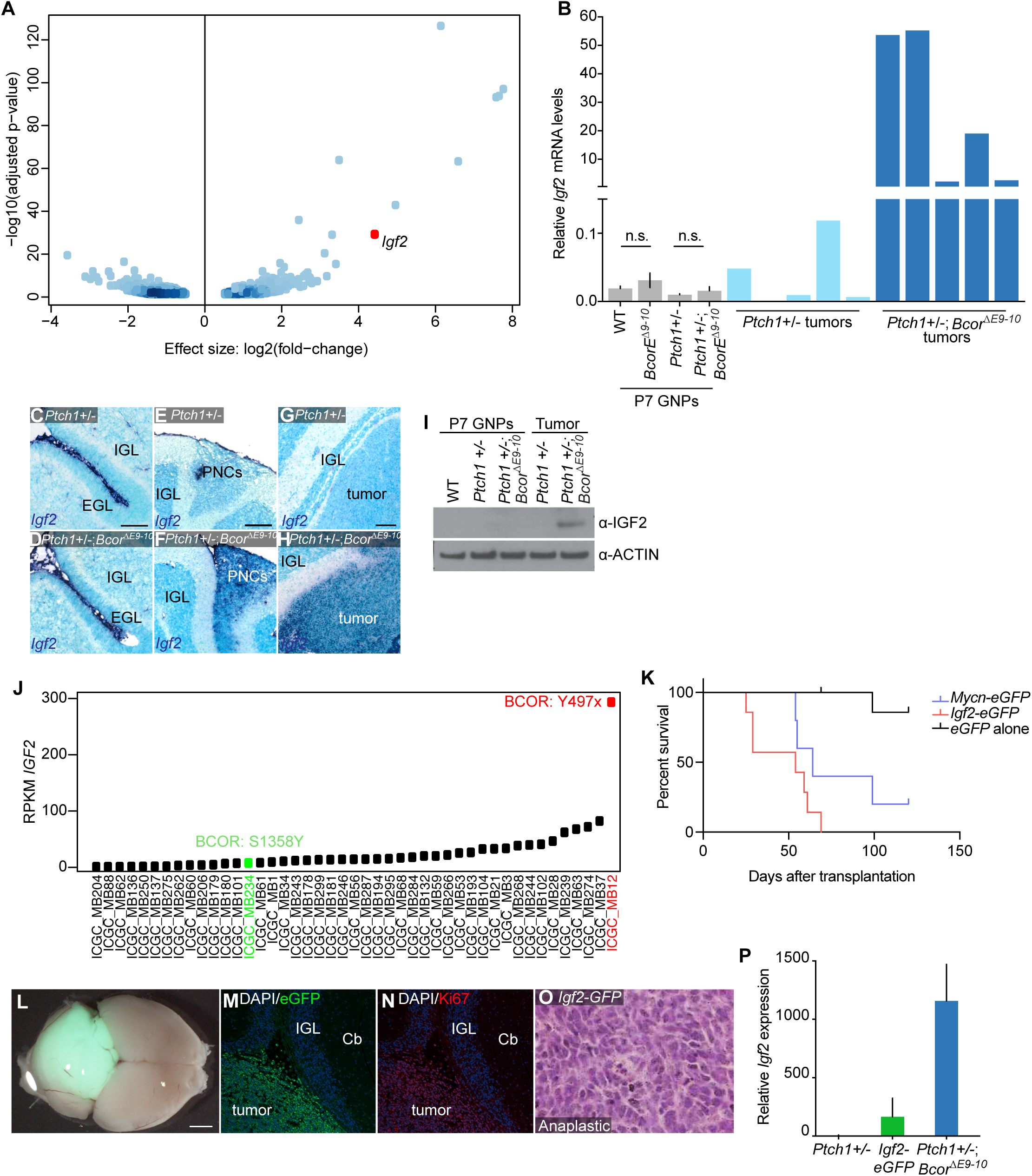
*Igf2* is overexpressed in *Ptch1*+/-;*Bcor*^*ΔE9-10*^ tumors. (**A**) Volcano plot of differentially expressed genes between *Ptch1*+/- vs. *Ptch1*+/-;*Bcor*^*ΔE9-10*^ tumors. (**B**) Relative mRNA expression of *Igf2* (quantitative PCR) of P7 GNPs (gray) in indicated genotypes, *Ptch1+/-* tumors (light blue), and *Ptch1*+/-;*Bcor*^*ΔE9-10*^ tumors (dark blue). n.s., not significant, Student’s t test. Bars of P7 GNPs represent mean +/- SEM. N = 6 (WT, *Bcor*^*ΔE9-10*^ GNPs) or N = 8 (*Ptch1*+/-, *Ptch1*+/-;*Bcor*^*ΔE9-10*^ GNPs). (**C-H**) *in situ* hybridization of antisense *Igf2* probe in cerebellum of animals from the indicated genotypes. IGL, inner granule layer. EGL, external granule layer. PNC, preneoplastic cells. (**C**,**D**) P7. Scale bar, 100 μm. (E,F) P28. Scale bar, 200 μm. (G,H) Adult. Scale bar, 100 μm. (**I**) Anti-IGF2 western blot of total protein extracts from P7 granule neuron progenitors (GNPs) or tumor samples from the indicated genotypes. Anti-ACTIN, loading control. (**J**) Reads Per Kilobase of transcript, per Million mapped reads (RPKM) of *IGF2* from SHH-medulloblastoma samples (Northcott et al. 2017). (**K**) Survival curve of immunodeficient animals transplanted with 8×10^5^ *Ptch1*+/- GNPs transduced with retrovirus overexpressing *eGFP* alone (black, N=8), *Igf2*-*eGFP* (red, N=7) or *Mycn*-*eGFP* (blue, N=5). (**L**) Representative example of resulting *Igf2-eGFP* tumor from (K). Scale bar, 2000μm. (**M-N**) eGFP (M, green) and Ki67 (N, red) immunohistochemistry of *Igf2-eGFP* driven tumor. IGL, inner granule layer. Cb, cerebellum. (**O**) Representative histology sample of *Igf2-eGFP* driven tumor. (**P**) Relative mRNA expression of *Igf2* in tumor cells from indicated genotypes, Bars represent mean + SEM. N = 7 (*Ptch1+/-*), N=4 (*Igf2-eGFP)* or N = 5 (*Ptch1*+/-;*Bcor*^*ΔE9-10*^). See also Supplemental Figure S2, Supplemental Tables S1, S2.

We also verified *Igf2* overexpression using Affymetrix microarray data from an additional cohort of samples (Supplemental Fig. S2B,C, Supplemental Table S2) and by performing qPCR on individual tumor samples (Fig. 3B). Consistent with previous results (Corcoran et al. 2008), we did not detect obvious *Igf2* expression in P7 GNPs in wild-type animals (Fig. 3B). There was also no aberrant expression of *Igf2* in P7 GNPs from either *Ptch1*+/- or *Ptch1*+/-; *Bcor*^*ΔE9-10*^ mice by qPCR (Fig. 3B) nor ISH (Fig. 3C,D). Additionally, we did not detect robust *Igf2* expression in *Ptch1*+/- preneoplastic lesions or in *Ptch1*+/- tumors (Fig. 3B,E,G). Instead, aberrant *Igf2* expression was observed only in preneoplastic lesions of *Ptch1+/-;Bcor*^*ΔE9-10*^ animals (Fig. 3F) and in *Ptch1+/-;Bcor*^*ΔE9-10*^ tumors (Fig. 3B,H), suggesting that inactivation of both alleles of *Ptch1* is necessary for BCOR-dependent *Igf2* regulation as well. We also confirmed that IGF2 protein is expressed in *Ptch1+/-; Bcor*^*ΔE9-10*^ tumors, but not in P7 GNPs nor *Ptch1*+/- tumors (Fig. 3I).

To determine whether *Igf2* upregulation is a conserved mechanism across species, we examined human medulloblastoma samples in which both genomic and transcriptomic data are available (Table 1, Northcott et al. 2017). Notably, we discovered that the only case of *IGF2* overexpression was co-incident with a *BCOR* frameshift mutation (Fig. 3J). The *BCOR* mutation in this sample was confirmed using RNA-seq alignment data, and *BCOR* is highly expressed in this sample compared to other SHH-medulloblastoma samples (Supplemental Fig. S2D). While we also had one sample with a BCOR missense mutation (S1358Y) that did not exhibit upregulation of *IGF2* (Fig. 3J), this mutation is located within the pre-Ankyrin repeats and may not negatively affect BCOR function (Fig. 1A). BCOR expression in this tumor is also reduced compared to the rest of the cohort (Supplemental Fig. S2D), and the mutation was detected in only 2% of the RNA-seq reads.

### Igf2 overexpression is sufficient to drive tumorigenesis in Ptch1+/- GNPs

To verify that *Igf2* overexpression is an oncogenic cofactor in SHH-medulloblastoma formation (Corcoran et al. 2008), we transduced *Ptch1*+/- GNPs *ex vivo* with retroviruses encoding *Igf2-*IRES-e*GFP, Mycn-*IRES-e*GFP* as a positive control, or *eGFP* alone as a negative control. We transplanted 1×10^6^ transduced cells of each condition into the cerebellum of adult immunocompromised mice. *Igf2*-transduced cells formed aggressive tumors that developed significantly faster than cells transduced with *eGFP* alone (Fig. 3K, 100% penetrance; median survival = 54 days for *Igf2*, p < 0.0001, Log-rank (Mantel-Cox) test). *Igf2*-derived tumor formation was cell-autonomous, as proliferating tumor cells (Ki67-positive) were eGFP-positive (Fig. 3L-N). Histologically, these tumors resembled *Ptch1*+/-;*Bcor*^*ΔE9-10*^ tumors and all displayed an anaplastic histology (Fig. 3O, N=3).

Because *Igf2* overexpression accelerated tumorigenesis in this model, we examined the level of *Igf2* necessary for tumor induction and compared it to the expression level in the genetically engineered mouse models. We observed lower levels of *Igf2* expression in the *Igf2-*IRES-e*GFP* induced tumors compared to *Ptch1+/-;Bcor*^*ΔE9-10*^ genetic tumors (Fig. 3P), suggesting the level of *Igf2* in *Ptch1+/-;Bcor*^*ΔE9-10*^ tumors is more than sufficient to drive tumorigenesis.

### BCOR^ΔE9-10^ fails to interact with the PRC1.1 catalytic subunit RING1B, and the PUFD domain of BCOR is required for Igf2 repression

The C-terminal domain of BCOR mediates its interaction with the PRC1.1 complex, a multi-protein complex that transfers ubiquitin to H2AK119Ub via the E3 ubiquitin ligase RING1B to repress target gene transcription (Wang et al. 2004; 2018). We tested whether the PRC1.1 complex is affected in *Ptch1+/-;Bcor*^*ΔE9-10*^ tumors. Similar to previous studies (Tara et al. 2018), we found that BCOR^ΔE9-10^ no longer binds RING1B via co-immunoprecipitation in *Ptch1+/-;Bcor*^*ΔE9-10*^ tumor samples, while full-length BCOR and RING1B co-immunoprecipitate in *Ptch1+/-* tumor samples (Fig. 4A).

**Figure 4.**
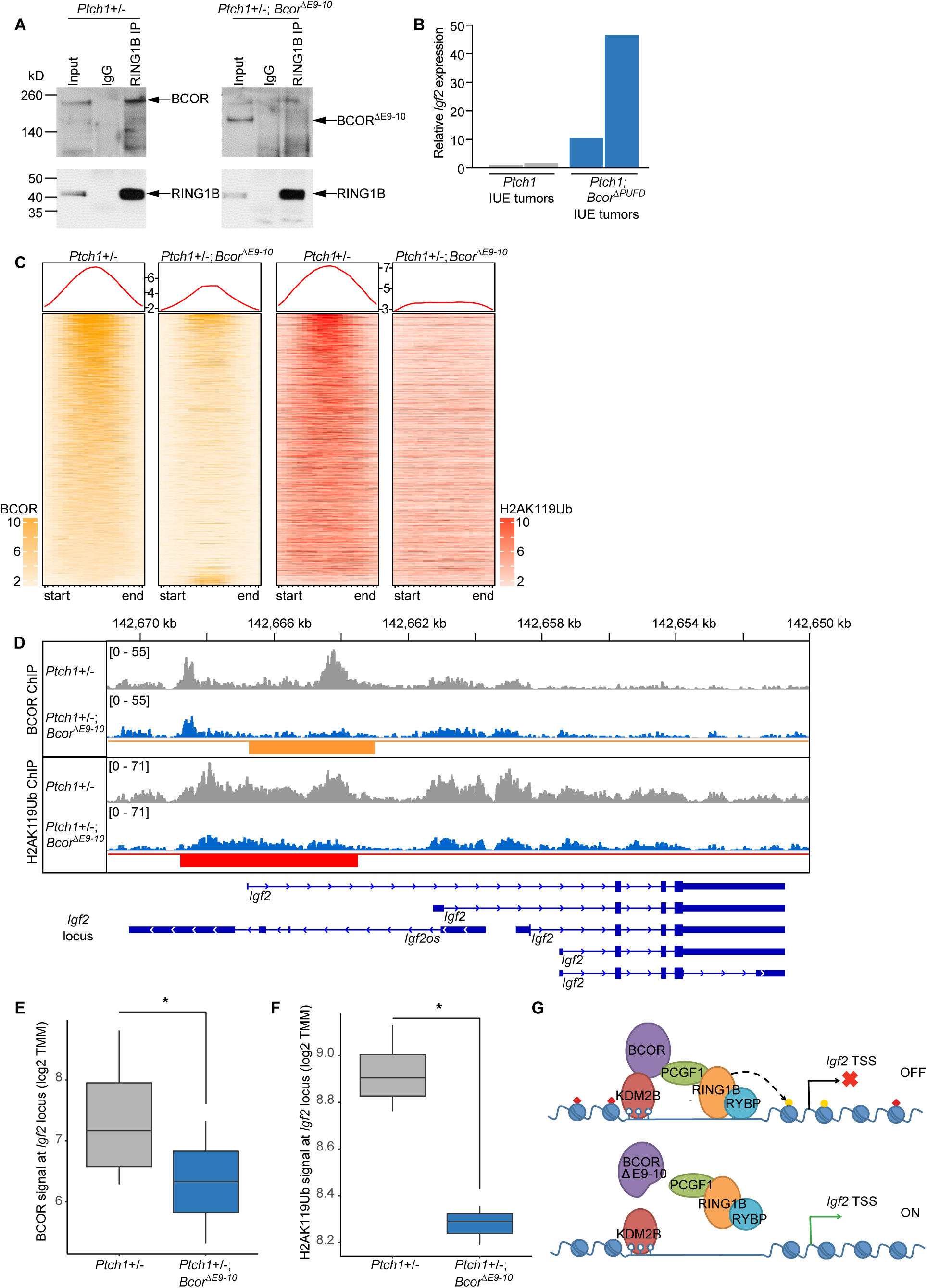
The PUFD domain of BCOR is required for *Igf2* repression through H2AK119 Ubiquitination. (**A**) Co-immunoprecipitation of BCOR or BCOR^ΔE9-10^ with RING1B in *Ptch1*+/- and *Ptch1*+/-;*Bcor*^*ΔE9-10*^ tumor cells, respectively. IgG, Immunoglobulin G control. (**B**) Relative mRNA expression of *Igf2* of two *Ptch1* CRISPR tumors (gray), and two *Ptch1;Bcor*^*ΔPUFD*^ CRISPR tumors (blue). (**C**) Heat map of significant differentially bound peaks from BCOR chromatin immunoprecipitation (ChIP) (yellow) between tumors from indicated genotypes and the status of H2AK119Ub ChIP signals (red) within these genomic loci. (**D**) Combined BCOR and H2AK119 ChIP peaks in indicated genotypes. Significantly different peak heights represented with yellow (BCOR) and red (H2AK119Ub) bars (BCOR peaks called first). *Igf2* locus represented in blue. (**E**) Quantification of BCOR ChIP signal at *Igf2* locus. Differential peak calling with DiffBind; *, p-adj <0.05. (**F**) Quantification of H2AK119Ub ChIP signal at *Igf2* locus. Differential peak calling with DiffBind; *, p-adj <0.05. (**G**) Model of PRC1.1 complex disruption. Open lollipops, unmethylated CpG. TSS, transcriptional start site. See also Supplemental Table S3, S4, Figure S3.

In the genetically engineered mouse model, BCOR^ΔE9-10^ is missing both the PUFD domain, which is responsible for PCGF1 and RING1B binding (Wang et al. 2018), and the C-terminal Ankyrin and Non-ankyrin repeats, which may interact with additional proteins outside of PRC1.1 (Fig. 1G), and therefore may also mediate tumor suppression independent of PRC1.1. To verify that loss of the PUFD domain alone, rather than the entire C-terminus, is responsible for *Igf2* activation during medulloblastoma formation, we conducted *in utero* electroporation (IUE) experiments to test the role of the PUFD domain specifically in tumorigenesis. We generated a single plasmid carrying tandem sgRNAs targeting *Ptch1* (Zuckermann et al. 2015) and *Bcor* at exon 13 (107 nucleotides upstream of the PUFD domain coding sequence in the linker region, Supplemental Table S3) and Cre recombinase. We injected this plasmid in the fourth ventricle of E13.5 embryos that carry a single copy transgene expressing a Rosa26::Lox-Stop-Lox-Cas9-IRES-GFP cassette (see Methods for details). In *Ptch1;Bcor*^*ΔPUFD*^ IUE tumors, *Igf2* is overexpressed compared to *Ptch1* IUE control tumors (Fig. 4B), suggesting that loss of BCOR PUFD (Table S3), likely via disruption of PRC1.1 complex assembly and targeting (Wang et al. 2018), aberrantly upregulates *Igf2* transcription.

### BCOR is bound at the Igf2 locus and represses gene transcription through H2AK119Ub

To determine whether BCOR is directly bound at the *Igf2* locus, we conducted BCOR chromatin immunoprecipitation (ChIP) in *Ptch1+*/*-* and *Ptch1*+/-;*Bcor*^*ΔE9-10*^ tumor samples (see Methods for details). The BCOR antibody we used was raised against a region common to BCOR and BCOR^ΔE9-10^ (encoded within exons 6-8) and thus detects both forms in Western blot (Fig. 1H) and ChIP. Global levels of BCOR binding are reduced but not completely lost in *Ptch1*+/-;*Bcor*^*ΔE9-10*^ tumors compared to *Ptch1+*/*-* tumors (yellow panels, 2063/2093 peaks, Fig. 4C, Supplemental Table S4). There is also a small set of genes with increased binding in the *Ptch1*+/-;*Bcor*^*ΔE9-10*^ samples (30/3093 peaks). Importantly, we also detect 3890 peaks that were unchanged between all the replicates of the two samples, suggesting that BCOR^ΔE9-10^ is indeed present. BCOR^ΔE9-10^ could be recruited by BCL6 to these loci but would be unable to interact with PRC1.1 components.

Next, we examined the *Igf2* promoter region in *Ptch1+/-* samples, and we found that indeed BCOR is associated with chromatin in that region (Fig. 4D). We then checked the same region in *Ptch1*+/-;*Bcor*^*ΔE9-10*^ and discovered the BCOR peak is significantly reduced at the locus (Fig. 4D,E, DiffBind, p-adj < 0.05). Given that BCOR^ΔE9-10^ no longer binds to the catalytic subunits of the PRC1.1 (Fig. 4A), residual BCOR^ΔE9-10^ would only be capable of PRC1.1-independent functions.

If BCOR^ΔE9-10^ renders the PRC1.1 non-functional, then H2AK119Ub levels should be reduced compared to *Ptch1*+/- controls. To test this hypothesis, we performed H2AK119Ub ChIP from chromatin prepared from the same tumor samples (see Methods for details). We found that total levels of H2AK119Ub decreased at peaks where BCOR signal is significantly decreased (red panels, Fig. 4C), indicating that the enzymatic activity of PRC1.1 complex is indeed reduced at loci with less BCOR. At the *Igf2* locus, H2AK119Ub levels are significantly reduced in *Ptch1*+/-;*Bcor*^*ΔE9-10*^ tumor samples (Fig. 4D,F). We also called differential H2AK119Ub peaks between the two tumor types and then examined BCOR binding status, observing a similar effect (Supplemental Fig. S3A).

In P7 cerebellum, BCL6 and BCOR are located at promoter regions of *Gli1* and *Gli2* (Tiberi et al. 2014). We examined these same regions in our samples, and we found decreased BCOR binding at these sites (Supplemental Fig. S3B,C). However, H2AK119Ub levels are not significantly different at these regions, and there is no corresponding overexpression of *Gli1* or *Gli2* in *Ptch1*+/-; *Bcor*^*ΔE9-10*^ tumors (Supplemental Fig. S3B,C Supplemental Table S1,S2). These results suggest that that the recruitment of BCOR by BCL6 is not required to maintain repression of *Gli1/Gli2.* The maintenance of H2AK119Ub at these loci suggests that other PRC1 complexes may compensate to prevent *Gli1/Gli2* overexpression.

Altogether, our results suggest that lack of full-length BCOR in *Ptch1*+/-;*Bcor*^*ΔE9-10*^ tumors leads to direct upregulation of *Igf2*, and this aberrant *Igf2* expression is closely associated with reduced H2AK119Ub and a disrupted PRC1.1 complex (Fig. 4G), enhancing the incidence of malignant transformation during tumor progression.

## DISCUSSION

### BCOR functions as a tumor suppressor in SHH-medulloblastoma, independent of BCL6

BCOR is a large, multi-domain protein implicated as a tumor suppressor in many pediatric cancers (Astolfi et al. 2019). In the case of medulloblastoma, *BCOR* mutations have been found together with mutations that result in aberrant activation of SHH signaling. To determine whether mutations in *BCOR* contribute to tumorigenesis, we used a *Ptch1* loss-of-function allele together with a conditional mouse *Bcor* allele, which mimics patient mutations. This allele, *Bcor*^*ΔE9-10*^, is similar to the human *BCOR* mutations, where a C-terminally truncated protein, retaining the BCL6 binding region but lacking the region for interaction with PRC1.1 components, is predicted to be produced. *Bcor*^*ΔE9-10*^ must be at least a partial loss-of-function because we have removed the domain critical domain for PRC1.1 interaction, and we find the levels of the truncated protein are significantly reduced in GNPs. However, we cannot exclude the possibility that the residual protein retains activity, but this activity would have to be independent of PRC1.1. It has been proposed previously that BCOR, in conjunction with BCL6 and SIRT1, represses *Gli1/Gl2* (Tiberi et al. 2014). Although binding of BCOR to the *Gli1/Gli2* loci is significantly reduced, *Gli1/Gli2* mRNA expression is normal in the *Bcor*^*ΔE9-10*^ mutants, indicating that the residual activity is sufficient to repress *Gli1/Gli2*. Thus, our results show that the clinically relevant *BCOR* mutations modeled here act independently of the previously described BCL6/BCOR/SIRT1 regulation of *Gli1/Gli2* (Tiberi et al. 2014). Furthermore, recurrent *BCL6* mutations have yet to be identified in SHH-medulloblastoma patient samples, suggesting that BCOR and its PRC1.1-related functions may be more relevant to human tumor formation.

### BCOR directly represses Igf2 via PRC1.1 complex-mediated H2AK119 ubiquitination

Our results suggest that BCOR directly regulates *Igf2* transcription via interaction with other components of the PRC1.1 complex, which deposits H2AK119Ub repressive histone marks at the *Igf2* promoter. While our ChIP studies were conducted in progressive mouse medulloblastomas, this mechanism likely occurs in preneoplastic lesions of *Ptch1+/-* animals as well (given *Igf2* upregulation in *Ptch1*+/-;*Bcor*^*ΔE9-10*^ cerebellum). 85% of *Ptch1+/-* animals have preneoplastic lesions at P21, but only 15%-20% of animals develop progressive medulloblastoma (Corcoran et al. 2008; Kessler et al. 2009), suggesting that BCOR-PRC1.1 may function in preneoplastic cells to prevent aberrant *Igf2* overexpression. This hypothesis is further strengthened by the fact that IGF2 is necessary for late stage SHH-medulloblastoma progression (Hahn et al. 2000; Corcoran et al. 2008). Nevertheless, no recurrent mutations of genes that encode components of PRC1.1 other than *BCOR* have been identified in SHH-medulloblastoma. Perhaps these mutations influence cell viability in the developing brain, potentially leading to cell death rather than tumorigenesis.

If *Igf2* upregulation is indeed required to push tumor-prone *Ptch1+/-* cells towards tumorigenesis, our results nicely complement these findings. Lack of a functioning BCOR-PRC1.1 complex upregulates *Igf2* in preneoplastic lesions at P28 in *Ptch1*+/- animals, which then pushes these cells towards fast-growing, lethal tumors. Few molecular mechanisms of transformation of preneoplastic lesions to malignant cells have been uncovered, and the present study unveiled *Igf2* overexpression in preneoplastic cells as a likely mechanism. This general mechanism explains why all *Ptch1*+/-;*Bcor*^*ΔE9-10*^ mice, which robustly express *Igf2* early-on in preneoplastic lesions, develop tumors with significantly reduced latency and dramatically increased penetrance compared to *Ptch1+/-* mutations alone.

Given that mutations in *Bcor* alone do not lead to medulloblastoma, cooperating mutations are required to drive tumorigenesis. Based on our data, we think that mutations in *Bcor* cooperate with hyperactive SHH signaling to upregulate *Igf2*, which explains why we do not detect *Igf2* upregulation at P7. Presumably at P7, some single cells may have acquired mutations in *Ptch1* in both alleles, but these single cells would be difficult to detect in ISH slices at P7. However, these single cells then form preneoplastic outgrowths, which lack *Ptch1* and only express *Bcor*^*ΔE9-10*^, and thus overexpress *Igf2*.

Of note, mutations in *BCOR* and other PRC1.1 complex components have been identified in other tumor types. For example, the PRC1.1 complex is also important in preventing T-cell acute lymphoblastic leukemia (T-ALL). Mice lacking the PUFD domain of BCOR (same floxed allele as present study) or the zinc finger domain of KDM2B both developed lethal T-ALL (Tara et al. 2018; Isshiki et al. 2019), suggesting a broader role for PRC1.1 as a tumor suppressor in cancer. Notably, *Bcor*^*ΔE9-10*^ on its own was enough to induce T-ALL, suggesting a different mechanism is likely involved.

### IGF2 overexpression may be a general feature of BCOR altered tumors

Internal tandem duplications in the C-terminus of *BCOR* in CNS-HGNET-BCOR tumors exhibit upregulation of *IGF2* (Vewinger et al. 2019), suggesting that suppression of *IGF2* expression may be a general feature of BCOR function across pediatric brain tumor entities. Importantly, CNS-HGNET-BCOR tumors exhibit aberrant activation of the SHH pathway (Paret et al. 2016), suggesting that both SHH pathway activation and *BCOR* aberrations cooperate to activate *IGF2*, similar to our *Ptch1*+/-;*Bcor*^*ΔE9-10*^ tumors.

Some pediatric sarcomas are associated with *BCOR* mutations, including Clear Cell Sarcoma of the Kidney and Uterine Sarcomas, both of which carry internal tandem duplications in exon 15, similar to CNS-HGNET-BCOR tumors (Ueno-Yokohata et al. 2015; Roy et al. 2015; Mariño-Enriquez et al. 2018). Rhabdomyosarcomas also frequently harbor *BCOR* alterations (Seki et al. 2015; Demellawy et al. 2017). Intriguingly, *IGF2* overexpression is associated with *Ptch1*-related rhabdomyosarcoma (Hahn et al. 2000). It will be interesting to determine whether there is any link between *BCOR* alterations and *IGF2* overexpression in sarcomas, especially in sarcomas with SHH pathway aberrations.

As we uncovered, *Ptch1*-mutated tumors and *Ptch1*+/-;*Bcor*^*ΔE9-10*^ tumors have different tumorigenic properties, including histology, penetrance, and latency. If the association holds true across tumor samples, patients with *BCOR* and SHH-related mutations may benefit from different treatment strategies compared to SHH-medulloblastomas lacking *BCOR* mutations. Our results also demonstrate the necessity in investigating the cooperating mutations for each subtype of medulloblastoma, and by extension, other tumor subtypes, to understand how these mutations contribute to malignant phenotypes.

## Author Contributions

Conceptualization, N.V.B., P.A.N. and D.K.; Methodology, L.M.K., K.O., N.V.B., J.C., D.K.; Software, K.O.; Formal Analysis, L.M.K., K.O., J.C., D.K.; Investigation, L.M.K., N.V.B., J.C., P.B.G.S., M.V., S.v.R., L.S., B.S., N.M., B.A.O., A.K., D.K.; Data Curation, K.O.; Writing – Original Draft, L.M.K; Writing – Editing, L.M.K., K.O., N.V.B., M.D.G., V.J.B., S.M.P., P.A.N., D.K.; Writing – Review, all authors; Funding Acquisition, D.K., S.M.P.; Resources, M.D.G., V.J.B., M.K., A.M., O.A., S.M.P., P.A.N.; Visualization, L.M.K., K.O., J.C., D.K.; Supervision, S.M.P., P.A.N., D.K.

## Acknowledgments

We thank the Genomics and Imaging core facilities at DKFZ. We thank Ana Banito, Mija Blattner-Johnson and David T.W. Jones for their scientific input and discussions. This work was supported by a DFG grant to D.K., a Heinrich F.C. Behr Stipendium to N.V.B., and NIH grants 5R01CA071540 and R01HD084459 to V.J.B.

---

**Supplemental Figure S1.**
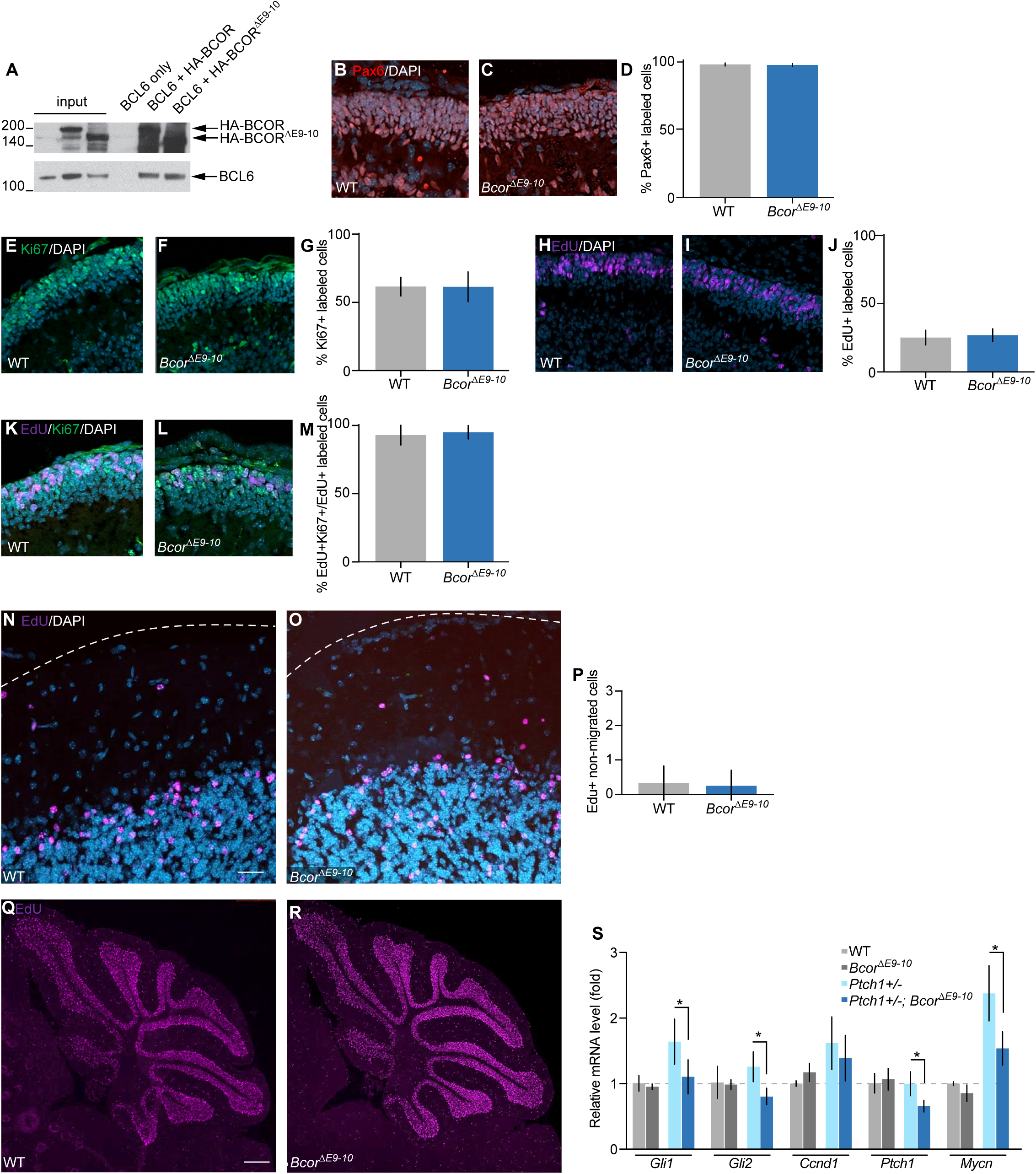
*Bcor*^*ΔE9-10*^ does not affect differentiation or migration of granule neuron progenitors in the developing cerebellum. Related to Figure 1. (**A**) Co-immunoprecipitation of HEK293T cell extract expressing HA-tagged full-length BCOR or HA-tagged BCOR^ΔE9-10^ with BCL6. 5% input. IP: anti-HA. Top blot: anti-HA. Bottom blot: anti-BCL6. (**B**) PAX6 (red) immunohistochemistry in P7 external granule layer in wild type (WT) animals. (**C**) Same as in B, except in *Bcor*^*ΔE9-10*^ animals. (**D**) Quantification of B,C. Bars represent mean +/- standard deviation. N = 12 regions. (**E**) P7 wild type animals were injected with a single dose of EdU (purple) and sacrificed 24h later to monitor proliferation of GNPs. (**F**) Same as in E, except in *Bcor*^*ΔE9-10*^ animals. (**G**) Quantification of E,F. Bars represent mean +/- standard deviation. N = 12 regions. (**H**) Ki67 (green) immunohistochemistry in P7 external granule layer in wild type (WT) animals. (**I**) Same as in H, except in *Bcor*^*ΔE9-10*^ animals. (**J**) Quantification of H,I. Bars represent mean +/- standard deviation. N = 12 regions. (**K**) P7 wild type animals were injected with a single dose of EdU (purple) and sacrificed 24h later to monitor proliferation (Ki67, green) of GNPs. (**L**) Same as in K, except in *Bcor*^*ΔE9-10*^ animals. (**M**) Quantification of K,L. Bars represent mean +/- standard deviation. N = 12 regions. (**N**) P7 wild type animals were injected with a single dose of EdU (purple) and sacrificed at P28 to monitor migration of GNPs. (**O**) Same as in N, except in *Bcor*^*ΔE9-10*^ animals. (**P**) Quantification of N,O. Bars represent mean +/- standard deviation. N = 12 regions. (**Q**) Same as in M, but EdU (purple) staining of entire cerebellum. (**R**) Same as in Q, except in *Bcor*^*ΔE9-10*^ animals. (**S**) Relative mRNA expression levels of indicated genes in P7 granule neuron progenitors from wild type (light gray), *Bcor*^*ΔE9-10*^ (dark gray), *Ptch1*+/- (light blue) and *Ptch1*+/-;*Bcor*^*ΔE9-10*^ (dark blue) animals. *, p<0.05, Student’s T test. Bars represent mean +/- standard deviation. N = 3 (WT, *Bcor*^*ΔE9-10*^ GNPs) or N = 4 (*Ptch1*+/-, *Ptch1*+/-;*Bcor*^*ΔE9-10*^ GNPs).

**Supplemental Figure S2.**
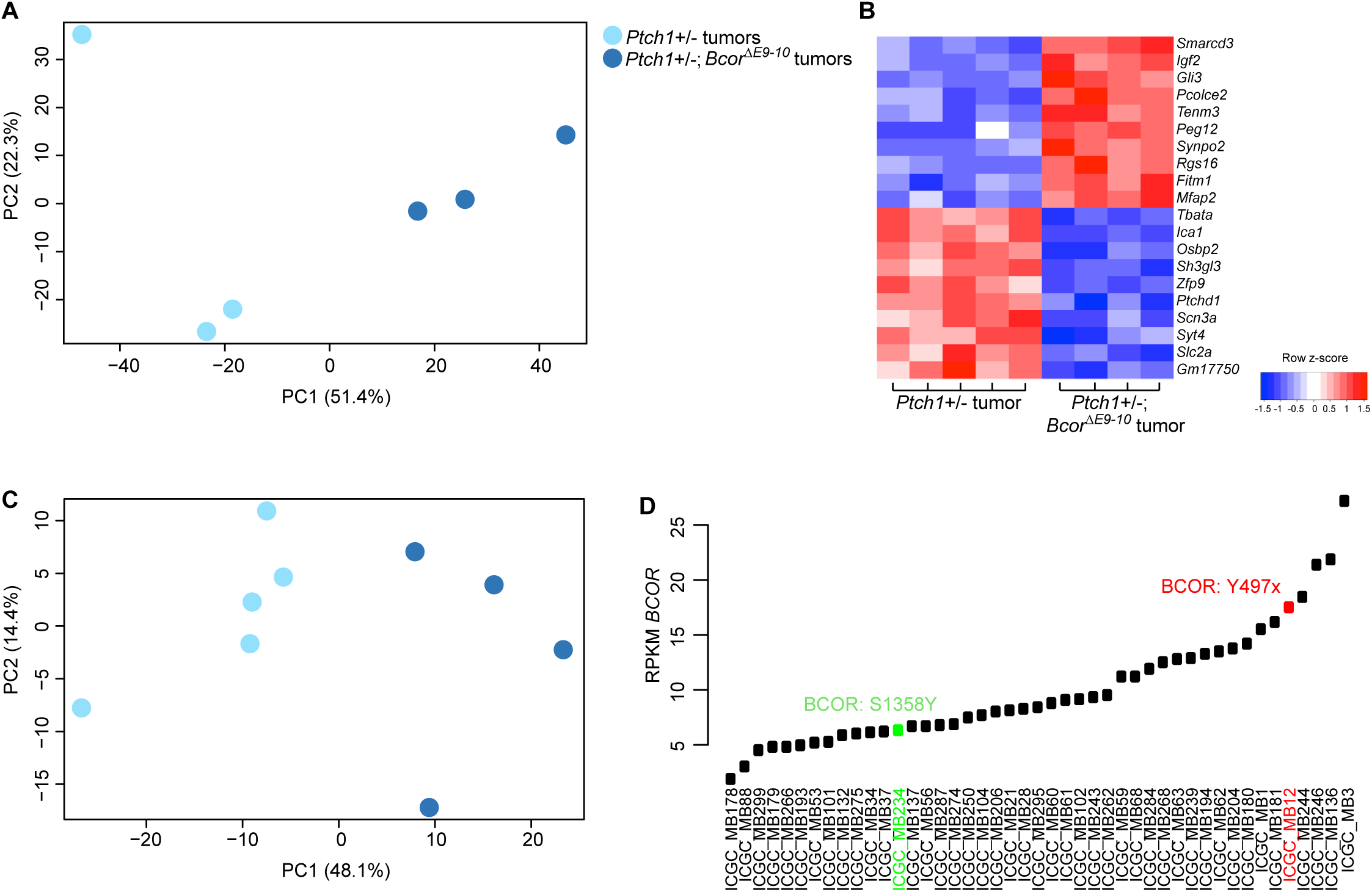
Further investigation of *Bcor*-mutated tumors. Related to Figure 3. (**A**) Principal component analysis of RNA-seq gene expression profiles from *Ptch1*+/- vs. *Ptch1*+/-;*Bcor*^*ΔE9-10*^ tumors using top 500 differentially expressed genes. (**B**) Heat map of top differentially expressed genes in *Ptch1*+/- vs. *Ptch1*+/-;*Bcor*^*ΔE9-10*^ tumors. (**C**) Similar to (A), but using Affymetrix gene expression profiles. (**D**) Reads Per Kilobase of transcript, per Million mapped reads (RPKM) of *BCOR* from SHH-medulloblastoma samples (Northcott et al. 2017).

**Supplemental Figure S3.**
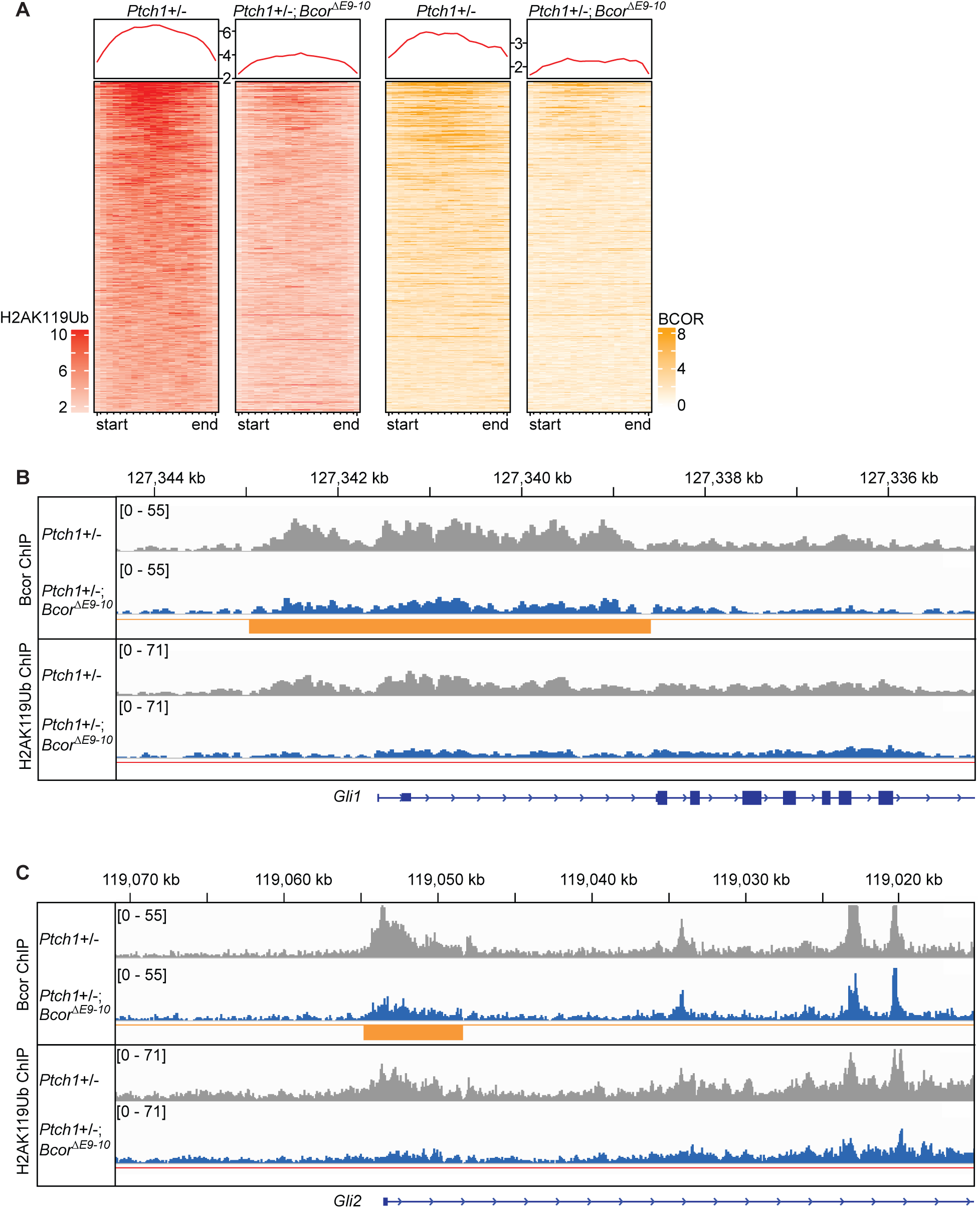
Status of BCOR binding at H2AK119Ub differential peaks and differential BCOR peaks at the *Gli1/ Gli2* loci. Related to Figure 4. (**A**) Heat map of significant differentially bound peaks from H2AK119Ub ChIP (red) between tumors from indicated genotypes and the status of BCOR ChIP peaks (yellow) within these genomic loci. (**B**) Combined BCOR and H2AK119 ChIP peaks in indicated genotypes at the *Gli1* locus. Significantly different peak heights represented with yellow (BCOR). (**C**) Same as in (B), but at the *Gli2* locus.

## Supplemental Table Legends

**Supplemental Table S1. Differentially expressed genes between *Ptch1*+/- vs. *Ptch1*+/-;*Bcor***^***ΔE9-10***^ **tumors.** Related to Figure 3. RNA-seq data, min log2FC 0.5, min. adjusted p-value 0.05 tumor_RNA_seq_Ptch1Het_BcorKO_vs_WT.qval_0.05.log2FC_0.5

**Supplemental Table S2. Differentially expressed genes between *Ptch1*+/- vs. *Ptch1*+/-;*Bcor***^***ΔE9-10***^ **tumors.** Related to Figure 3. Affymetrix data, top 1000 most confident genes based on sorted adjusted p-value. tumour_BcorWT_vs_BcorKO_top1000DEG.18042017.

**Supplemental Table S3. *Bcor*-PUFD mutations in *in utero* electroporation-derived tumors.** Related to Figure 4. *Bcor*-PUFD sgRNA and Sanger sequencing at PAM site with predicted protein change.

**Supplemental Table S4. Quality control overview and differentially binding peaks between *Ptch1*+/- vs. *Ptch1*+/-;*Bcor***^***ΔE9-10***^ **tumors across BCOR and H2AK119ub ChIP signals.** Related to Figure 4. Each differential peak is annotated with assigned closest gene, min adjusted p-value 0.05.

**Supplemental Table S5. Oligo sequences for genotyping and qPCR.** Related to Materials and Methods.

## Materials and Methods

### Animals

*Bcor* conditional knockout mice were generated by breeding of *Bcor*^*fl/fl*^ (*loxP* sites flanking exons 9 and 10, Hamline *et al. in prep*.) with *Atoh1-Cre (*JAX, #011104) and *Ptch1* heterozygous mice (Goodrich et al. 1997). Genotyping primer sequences are available in Supplemental Table S5. All mice were bred in a C57BL/6N background. For tumor studies, only homozygous *Bcor*^*fl/fl*^ females or *Bcor*^*fl/y*^ males were used and referred as *Bcor*^*ΔE9-10*^. Of note, *Bcor*^*fl/+*^ females developed medulloblastoma at similar rates. For the *in utero* electroporation experiments, male *Rosa26-CAG-LSL-Cas9-P2A-EGFP* (JAX #024857) were crossed to CD-1 female animals purchased from Janvier and checked daily for vaginal plug. For transplantation experiments, Nod-scid IL2Rgamma^null^ (NSG, JAX #005557) were used. All animal experiments for this study were conducted according to the animal welfare regulations approved by the Animal Care and Use Committee of the National Institute of Neuroscience, NCNP, Japan (approval number 2019028R2), and the responsible authorities in Baden-Württemberg, Germany (Regierungspraesidium Karlsruhe, approval numbers: G-182/13, G-48/14, G-64/14, G-29/18, and G-23/19).

#### in utero electroporation

*in utero* electroporation was performed as previously described (Feng et al. 2017). For these experiments, we used a single px330 plasmid (Addgene #42230) modified to carry two sgRNA cassettes, and Cre recombinase coding sequence was used in place of Cas9. This plasmid together with a luciferase encoding plasmid was injected into the fourth ventricle of E13.5 embryos and electroporated (30V, 50 ms-on, 950 ms-off, 5 pulses). Positive pups were identified by luciferase signal at P7 and animals were monitored daily for the first neurological signs of medulloblastoma formation.

#### Orthotopic transplantation

For retransplantation experiments, 8×10^5^ freshly isolated, purified tumor cells from *Ptch1*+/- or *Ptch1+/-;Bcor*^*ΔE9-10*^ animals were retransplanted to the cerebellum of NSG mice, according to published SOP (Brabetz et al. 2018), and animals were monitored daily for signs of tumor growth. For *Igf2* overexpression studies, retroviruses were freshly generated as previously described (Kawauchi et al. 2012). 1×10^6^ P7 GNPs isolated from *Ptch1+/-* mice were cultured *in vitro* and transduced with MSCV-based retroviruses carrying *Igf2-IRES-eGFP, Mycn-IRES-eGFP* (positive control), or *eGFP* only (negative control). Two days later, GFP+ GNPs were harvested and transplanted to NSG mice in the same manner as above.

#### EdU injections and imaging

P7 mice of the appropriate genotype were injected intraperitoneally with 10 mg/ml stock solution of EdU (50 mg/kg, 20ul total volume, dissolved in PBS, Invitrogen). Mice were sacrificed at P8 (proliferation) or P28 (migration). EdU labeling reaction was conducted as described in (Zeng et al. 2010). Brains were harvested, fixed overnight in 4% paraformaldehyde (PFA) at 4°C, soaked in 30% sucrose in PBS overnight, and embedded in Optimal Cutting Temperature (OCT) compound. Frozen 10μm-thick sections were collected on Fisher Superfrost Plus slides using a cryostat and treated with Click-iT™ EdU imaging reagent (Invitrogen). For quantification, we counted the number of average EdU positive cells in four 100 μm x 100 μm regions in 4 independent cerebellum.

### Plasmids

A single px330 plasmid modified to carry Cre instead of Cas9 was generated to carry tandem cassettes for sgRNAs against *Ptch1* (CTGGCCGGAAAGCGCCGCTG) and *Bcor* (ATAGAACTCCCAAGCGCCGC), using ASAP-cloning protocol (Zuckermann et al. 2018). For *Ptch1* only targeting construct, the sgRNA for *Bcor* was replaced with a control sgRNA (GCGACCAATACGCGAACGTC). For MSCV-based plasmids, the coding region for *Igf2, Mycn*, or empty vector was cloned using BamHI and XhoI sites. For co-IP plasmids, full-length *Bcor* cDNA or truncated *Bcor*^*ΔE9-10*^ cDNA was cloned with an N-terminal HA tag into pCAG vector using InFusion (Takara). pCMV-SPORT 6.1 with mouse *Bcl6* cDNA was purchased from Horizon Discovery (Clone ID 6309948).

### *in situ* hybridization

*in situ* hybridization (ISH) was performed as described previously (Kawauchi et al. 2006). Plasmids containing the coding region of *Bcor* (Clone ID 6412868, Horizon Discovery), *Igf2* (Clone ID 30013295, Horizon Discovery), and *Ccnd2* (5716186, Horizon Discovery) were linearized with XhoI, EcoRI, and AccI, respectively, and used to generate antisense RNA probes (DIG RNA labeling kit, Roche). Probes were hybridized on 10μm-thick cryosections of the cerebellum or tumor samples of interest and counterstained with Methyl green (Sigma). For expression pattern analysis, we checked three independent cerebella and all ISH results reported in this paper were consistent in all three samples.

### Histopathology and Immunostaining

For histopathology, samples of mouse medulloblastomas (3 tumors per genotype) were formalin-fixed, paraffin-embedded and sectioned at 5 μm. The sections were stained with hematoxylin and eosin (H&E), and histological classifications were performed blinded to genotype.

Immunostaining was performed as previously described (Pajtler et al. 2019). Brains were harvested, fixed overnight in 4% paraformaldehyde (PFA) at 4°C, soaked in 30% sucrose in PBS overnight, and embedded in Optimal Cutting Temperature (OCT) compound. Frozen 10μm-thick sections were collected on Fisher Superfrost Plus slides using a cryostat. Sections were blocked for 30min at RT with 10% normal donkey serum (NDS) and incubated with primary antibody overnight at 4°C. After washing with PBST, sections were incubated with appropriate fluorescence secondary antibodies for 1h at RT. Slides were mounted in ProLong Gold Mountant (Invitrogen). Antibodies used were anti-Ki67 (ab15580, abcam, 1:500), anti-GFP (ab13970, abcam, 1:1000), anti-PAX6 (PPRB-278P, Covance, 1:500). All slides were counterstained with DAPI (300 nM). For quantification, we counted the number of average positive cells in four 100 μm x 100 μm regions in 3 independent cerebellum.

### Quantitative PCR (qPCR)

GNP isolation was performed as previously described (Kawauchi et al. 2012). For *in vitro* studies, 5 million cells were plated per well in a 6-well plate and cultured for 48h in the presence or absence of 200 nM smoothened agonist (Merck). RNA was extracted and cDNA was generated with Superscript II kit (Invitrogen), according to manufacturer’s instructions. RNA and cDNA from tumor samples was generated similarly. Relative gene expression was compared to *Gapdh* in all experiments except Fig. 3P and 4B, which was compared to the geometric mean of *Hrpt1, Rpl27*, and *Rer1* (Thomas et al. 2014). All primer sequences are listed in Supplemental Table S5.

### Western blotting

Protein was extracted from tissue or cells by addition of RIPA Buffer (Sigma Aldrich) including complete Protease Inhibitor Cocktail Tablets (Roche) according to manufacturer’s instruction. Tissue was homogenized by using the Tissue Master 125 Homogenizer (OMNI International) or sonication after adding lysis buffer directly to the frozen sample. Cell lysates were vortexed 3-4 times every 10 min for a total of 30 min and were kept on ice in between. Then lysates were centrifuged at 17000*g* for 30 min at 4°C. Protein concentration of cell lysates was measured using Pierce BCA Protein Assay (Thermo Fisher Scientific), following supplier’s instructions. Primary antibodies include anti-BCOR (rabbit polyclonal, Vivian Bardwell Lab; University of Minnesota Cat# anti-BCOR, RRID:AB_2716801, 1:1000), anti-ACTIN-HRP (ab49900, abcam, 1:25000), anti-BCL6 (#4242, CST, 1:1000), anti-RING1B (#39663, Active Motif, 1:500), anti-HA (#MMS-101P, BioLegend, 1:1000).

### Co-immunoprecipitation

Co-immunoprecipitation studies of BCOR and BCL6 were performed similar to (Huynh et al. 2000). Briefly, we overexpressed cDNAs expressing HA-tagged full-length BCOR or BCOR^ΔE9-10^ with BCL6. We also included a BCL6-only negative control. We used HEK293T cells for these experiments (CRL-3216). Three days after transfection, we harvested the cells and extracted protein in LB200 buffer (20 mM Tris, pH 8, 1% Triton-X, 1 mM EDTA, 1mM EGTA, 200 mM NaCl, Complete protease inhibitors (Roche)) with sonication. After preclearing anti-HA beads (Pierce) with 1% BSA in PBS, 51 μg of total protein lysate were bound to the beads for 1h at 4°C, rotating. After three wash steps with LB200 at 4°C, we eluted the bound proteins with 2x SDS Laemmli buffer and ran total lysate on 4-12% bis-Tris gradient gel (NuPAGE, Thermo Fisher Scientific) for downstream Western blotting.

For interaction tests between RING1B and BCOR-FL / BCOR^ΔE9-10^ in mouse tumor samples, tumor cells were lysed in LB200 buffer with sonication. Anti-RING1B antibody was bound to IgG beads (#39663, Active Motif, 1:500) for 1h at RT before applying 500 μg of the total cell lysate. Proteins were allowed to bind for 4h at RT, rotating. After three wash steps with LB200 at RT, bound proteins were eluted as above.

### X-gal whole mount staining

X-gal staining was performed as previously described (Zindy et al. 2014). Brains were fixed in fresh 4% PFA / PBS for 1 hour on ice, rinsed in x-gal Rinse Buffer (100 mM sodium phosphate, pH 7.3, 2 mM MgCl_2_, 0.01% sodium deoxycholate, 0.02% (w/v) NP-40) 3×10min, followed by x-gal Staining Solution (5 mM potassium ferricyanide, 5 mM potassium ferrocyanide, 1 mg/ml X-gal (in DMF)) 37°C overnight. After, brains were placed in formalin (10%) and the number of preneoplastic lesions per cerebellum was quantified on a stereomicroscope (N=3 per genotype).

### Human tumor data analysis

The presence of *BCOR* mutations and indels was integrated from the medulloblastoma landscape study (Northcott et al. 2017). For available samples, the *IGF2* gene expression was checked from RNA-sequencing data (Fig. 3J). Additionally, the *BCOR* somatic indel in sample MB12 was confirmed via RNA-seq alignment data. We only detected 2% of reads with the missense mutation in MB234, however.

### Mouse model gene expression analysis

#### RNA extraction

Tissue was homogenized using the Tissue Master 125 Homogenizer (OMNI International) and/or QIAshredder (Qiagen). RNA was extracted with RNeasy Mini Kit (Qiagen) following supplier’s instructions. For measurement of RNA quantity, ND1000 Spectrophotometer (NanoDrop Technologies) and/or Agilent 2100 Bioanalyzer was used. RNA sequencing libraries were processed and sequenced on a HiSeq V4 (SR 50) by the DKFZ Genomics Core Facility.

#### RNA-seq analysis

Initial transcriptome profiling was performed using RNA-sequencing data from *Ptch1*+/- (N=3) and *Ptch1*+/-;*Bcor*^*ΔE9-10*^ (N=3) medulloblastoma tumor models. The alignment was performed using STAR v2.4.1 tool (Dobin et al. 2013) to mm10 reference genome, and gene expression counts were computed using featureCounts module of the Subread package v1.4.6 (Liao et al. 2014) with Ensembl GRCm38 v72 annotation. The quality control was performed with Qualimap v2 mode RNA-seq QC (Okonechnikov et al., 2016) and differentially expressed genes were detected from application of DESeq2 R package (Love et al. 2014) with applied limits: min adjusted p-value 0.05, min log 2FC 0.5.

Additional extended transcriptome profiling from *Ptch1+/-* (N=5) and *Ptch1*+/-;*Bcor*^*ΔE9-10*^ (N=4) medulloblastoma SHH tumor models was performed with Affymetrix microarray 430v2. General quality control and normalization were performed using Affy package (Gautier et al. 2004) followed by principal component analysis and unsupervised hierarchical clustering. The differentially expressed genes were detected using Limma package (Ritchie et al. 2015) and sorted using adjusted p-value limit for the top 1000 selection.

#### ChIP-sequencing data analysis

Chromatin Immunoprecipitation of H2AK119Ub (Cell Signaling Technologies) and BCOR (Vivian Bardwell Lab; University of Minnesota Cat# anti-BCOR, RRID:AB_2716801) for *Ptch1*+/-;*Bcor*^*ΔE9-10*^ and *Ptch1+/-* tumor samples was performed by Active Motif. One sample of *Ptch1*+/-;*Bcor*^*ΔE9-10*^ was discarded for outlier levels of *Igf2* revealed by RNA-seq after ChIP analysis. The sequencing reads representing H2AK119Ub and BCOR obtained from *Ptch1+/-* (N=4) and *Ptch1*+/-;*Bcor*^*ΔE9-10*^ (N=3) tumor model samples were aligned to the mm10 and Drosophila spike-in references using bwa 0.6.2 (Li and Durbin 2009). Next the general quality control and read alignments statistics collection was performed using Qualimap v2.2 toolkit mode BAM QC (Okonechnikov et al. 2016). The coefficients from Drosophila spike-ins were computed based on the alignment statistics and applied to adjust to the reads distribution per sample as suggested by Active Motif. For peak calling Macs 1.4 tool (Zhang et al. 2008) was applied with corresponding background included as input and using p-value limit 1e-9. Differential peaks were detected using DiffBind R package (Ross-Innes et al. 2012) with min adjusted p-value limit 0.05.

